# Temporal integration in the subcortical auditory system and behavioral evidence of its dysfunction after “temporary” noise-induced hearing loss

**DOI:** 10.64898/2026.07.14.738495

**Authors:** Chase A. Mackey, Jane A. Mondul, Ramnarayan Ramachandran

**Author notes:** The authors declare no conflicts of interest. Email addresses: Chase Mackey Jane Mondul Ramnarayan Ramachandran.

## Abstract

How sensory information is processed over time is often conceptualized as a process of temporal integration. Recently, auditory temporal integration has received renewed attention as a potential assay of hidden hearing loss caused by cochlear synaptopathy in rodent and avian studies. How these results relate to human hearing is in question due to a lack of studies in primates, and, more generally, the neural basis of auditory temporal integration is unclear, as most subcortical studies of it have been conducted under anesthesia. We have recently introduced a nonhuman primate (NHP) model which can address translational questions about auditory temporal integration and hidden hearing loss. Thus, in this study, we utilized single-unit recordings and compared derived neurometric measures to psychometric measures of temporal integration in normal hearing NHPs performing a tone-in-noise detection task. We then assessed psychometric measures of temporal integration in NHPs before and after noise exposure. In normal hearing NHPs, cochlear nucleus and inferior colliculus (IC) integration rates were significantly greater than psychometric rates. However, in noise only, ∼25% of IC neurons exhibited similar integration rates to behavior. After noise exposure, psychometric integration was disrupted for brief stimuli presented in quiet, but not in noise. The dynamic range of the psychometric function reliably increased, months after recovery from the noise-induced temporary threshold shift (TTS). Together, these data identify a subcortical neural substrate for temporal integration in noisy environments and suggest that behavioral assays of temporal integration may serve as sensitive indicators of subclinical hearing loss.

## INTRODUCTION

The process by which sensory evidence is summed up over time is characterized by a variety of different models across sensory systems (Heil et al., 2017; Hernández-Pérez et al., 2020; Huk and Shadlen, 2005; Liu et al., 2015; Viemeister and Wakefield, 1991; Watson, 1979). These models of temporal integration account for a wide variety of behavioral data, and their explanatory success emphasizes the importance of relying on converging lines of evidence from modeling, psychophysics, and neurophysiology to characterize perceptual processes. Within the auditory domain, the *rate* at which threshold changes with duration is typically used to quantify integration (Costalupes, 1983; O’Connor et al., 1999; Watson and Gengel, 1969), and the clinical utility of temporal integration has been well characterized in studies of short-tone audiometry (Sanders and Honig, 1967).

Neurophysiological investigations of auditory temporal integration have differed in their conclusions regarding where neuronal integration rates match behavioral integration rates. (Heil et al., 2008) leveraged anesthetized cat auditory nerve recordings to suggest temporal integration is essentially accomplished in the periphery. Studies in anesthetized chinchillas suggest otherwise: integration rates derived from the cochlear nucleus (CN) approximate psychophysical measures better than the auditory nerve (Clock et al., 1993; Clock Eddins et al., 1998; Viemeister et al., 1992). Other studies have suggested primary, or higher-order, auditory cortex to match behavioral integration rates (Lütkenhöner, 2011) (JUFANG, 1997). And one of the few causal studies of auditory temporal integration implicates auditory cortical projections to parietal cortex (Yao et al., 2020). Finally, the superior colliculus may represent a neural integrator of auditory information as in the visual domain (Jun et al., 2021), and as studies of sensorimotor integration suggest (Jay and Sparks, 1987; Rajala et al., 2017; Wallace et al., 1996). Despite these compelling possibilities, it is difficult to compare subcortical and cortical studies of auditory temporal integration. Subcortical studies have generally utilized anesthetized animals, but task-engagement has effects at least as early as the inferior colliculus (IC) (Rocchi and Ramachandran, 2020; Shaheen et al., 2021; Slee and David, 2015). Here we compared psychometric and neurometric measures of temporal integration derived from single-unit responses in the CN and IC of behaving macaques to further evaluate our hypothesis that the IC better correlates with perceptual measures due to neuronal convergence (Mackey et al., 2024, 2023).

Motivation for the second part of this study stems from the fact that many individuals with normal hearing thresholds report auditory perceptual deficits and display large individual differences in complex psychoacoustic tasks (Parthasarathy et al., 2020; Ruggles et al., 2011; Tremblay et al., 2015). Cochlear synaptopathy (the selective dysfunction of inner hair cell synapses) is a potential mechanisms underlying such difficulties and is a promising mechanistic target for enhancing the diagnosis and treatment of such deficits, given its neurophysiological effects on stimulus encoding (Asokan et al., 2018; Furman et al., 2013; Kujawa and Liberman, 2009; Shaheen et al., 2015; Shaheen and Liberman, 2018; Suthakar and Liberman, 2021).

Specifically, noise-induced temporary threshold shifts (TTS) can induce cochlear synaptopathy, reducing auditory nerve population responses as indexed by the auditory brainstem response (ABR), without altering ABR thresholds. Theoretically this could lead to perceptual deficits, such as impaired temporal integration, due to altered release dynamics and signal propagation (e.g. Kohrman et al., 2020; Moverman et al., 2023; Pal et al., 2025; Sheets et al., 2017; Wan and Corfas, 2017). However, pharmacological analogs of noise-induced cochlear synaptopathy in studies in budgerigars and chinchillas have not produced such deficits (Trevino et al., 2022; Wong et al., 2019). Here, we re-evaluate this possibility in the context of our recently introduced nonhuman primate (NHP) model of noise-induced hidden hearing loss following TTS, within which we have noted perceptual deficits, neural deficits, and altered synaptic morphology following recovery from temporary threshold shifts (Mackey et al., 2026; Mondul et al., 2026). In this model, NHPs display hypertrophied presynaptic ribbons, suggesting altered afferent vesicular pools / release and associated neural signaling, impaired spatial processing and reduced binaural interaction in the ABR, potentially indicating a reduction in the temporal integration necessary for binaural circuit function (Mackey et al., 2026). Noise-induced TTS could thus alter temporal integration more generally, which would have a wide range of implications beyond studies of spatial hearing. This led us in the present study to test the hypothesis that psychophysical measures of temporal integration would be degraded by noise exposure that causes TTS but does not permanently elevate audiometric thresholds or cause hair cell loss.

## METHODS

### Subjects

Seven adult male rhesus macaques (*Macaca mulatta*) took part in the study. Their ages were 5 – 10 years. All weighed 10 – 12 kg. Macaques were implanted with headposts (all) and recording chambers (A, Br, C, D). Monkeys A, Br, C, and D participated in the simultaneous neural and behavioral recordings (the detection task described below using 200 ms tones). Monkeys Bi, Ha, and Ga participated in the behavioral experiments and noise exposure described below, where tones of different durations were used. Four adult female rhesus macaques (Lu, Ne, Op and Pi) participated in the behavioral experiments before and after noise exposure. At the beginning of the study, ages ranged from 6-8 years old, and body weights ranged from 5.5-7 kg. Macaques were fed LabDiet Monkey Diet 5037 and 5050 (Purina, St Louis, MO). Macaques were provided produce and foraging toys. Environmental enrichment stimulated the auditory, visual, and olfactory systems on a rotational basis. Fluid consumption was managed: macaques received filtered municipal water averaging at least 20 ml/kg of body weight/day, but was often greater than or equal to 25 ml/kg/day. Weight was monitored at least weekly (typically 4 – 5 days each week) and stayed within bounds of the reference weights set to index the animal’s health while on study. A 12:12-h light:dark cycle was maintained and all procedures occurred between 8 AM and 6 PM during the light cycle. Visual, auditory, and olfactory contact with conspecifics was maintained within the housing room. The housing room was located in an AAALAC-accredited facility in accordance with the *Guide for the Care and Use of Laboratory Animals,* the Public Health Service Policy on Humane Care and Use of Laboratory Animals, and the Animal Welfare Act and Regulations. Macaques in this colony received routine health assessments and tuberculosis testing twice yearly. All research procedures were part of protocols that were approved by the Institutional Animal Care and Use Committee at Vanderbilt University Medical Center (VUMC).

### Surgical procedures

Monkeys were prepared for chronic experiments using standard techniques employed in nonhuman primate studies, and as reported in our previous studies (Dylla et al., 2013; Mackey et al., 2023). Briefly, anesthesia was induced via administration of ketamine and midazolam, and maintained via isoflurane. A metal headpost was implanted on the skull in order to restrict head movement during head fixation, minimizing sound pressure level changes at the ear as a result of positioning across behavioral sessions. The headpost was secured to bone using 8mm titanium screws (Veterinary Orthopedic Instruments) and encapsulated in bone cement (Zimmer Biomet). Circular chambers (Crist Instruments, Hagerstown, MD) were implanted over craniotomies to provide access to the cochlear nuclei and inferior colliculi (details in Mackey et al. 2023). Multimodal analgesics (pre- and post-procedure), intra-procedure fluids, and antibiotics (intra-procedure) were administered to the monkeys under veterinary oversight. Monkeys were monitored under veterinary supervision for at least 3 days post-surgery, until they were released for study.

### Stimuli

Tones and noise were generated using a sampling rate of 97.6 kHz. Tone frequencies were 0.5, 1, 2, 2.828, 4, 5.656, 8, 16, 24 and 32 kHz for the behavioral experiments, and could be 0.1-32 kHz in the neurophysiological experiments. Tone durations were defined as the entire tone waveform, including its rise/fall time. Tone durations were 3.25, 6.5, 12.5, 25, 50, 100, and 200 ms. Linear rise/fall times for tones were 1.25 ms for 3.25 ms tones, 2.5 ms for 6.5 ms tones, 4 ms for 12.5 and 25 ms tones, and 10 ms for the longer tone durations. Noise was broadband (5-40,000 Hz), 76 dB SPL (44 dB SPL for the single-unit experiments), and presented continuously from the same loudspeaker as the tones. For the simultaneous behavioral and neurophysiological experiments, tones were 200 ms in duration, of varying sound level (see **Behavioral Apparatus**), with 10 ms linear gate times.

### Behavioral apparatus

The behavioral apparatus has been described previously (Dylla et al., 2013; Mackey et al., 2023). Briefly, monkeys were seated in a custom designed primate chair situated inside a sound treated booth (IAC, model 1200A and Acoustic Systems model ER 247). Sounds were delivered from a free-field speaker (Rhyme Acoustics NuScale 216) located in the frontal field at a distance of 90 cm from the center of the monkey’s head. The speakers were calibrated with a ¼ inch microphone (378C01, PCB Piezotronics) that was located just outside the monkey’s ear canal. Care was taken to make sure that the speaker outputs were within 3 dB across all. Experimental flow was controlled by a computer running OpenEx software (System 3, TDT Inc., Alachua, FL). Macaques performed a lever-based reaction time Go/No Go tone detection task. Macaques initiated a trial by pulling a lever. Trials could be signal trials (80%) in which a tone signal of fixed duration was played after a random delay period of 800-3500 ms after the lever was pulled, or they could be catch trials (20%), in which no tone signal was played. The macaque was required to release the lever within a response window (600 ms after tone onset) to indicate detection on signal trials, and was required to continue to hold the lever on catch trials. Lever releases on signal trials (hits) were rewarded with fluid. Lack of lever release within 600 ms of the onset of the tone on signal trials (misses) was taken to indicate non-detection and was not rewarded or punished. Lever release on catch trials (false alarm) resulted in a 6-10 s timeout in which no trial could be initiated. Correct rejections (lack of release on catch trials) were not rewarded. Experiments were blocked by tone frequency and duration, while tone level was varied trial by trial, using the method of constant stimuli. Lever state was sampled at a rate of 24.4 kHz, leading to a temporal resolution of about 40 μs on the lever release.

### Noise exposure

The noise exposure protocol has been previously described in detail (Mondul et al., 2026). Monkeys were administered atropine, ketamine (10 mg/kg) and midazolam (0.05 mg/kg). Anesthesia was maintained with isoflurane (1.5 – 2%). The exposure noise was delivered in a closed-field system via MF1 speakers (TDT Inc.). The stimulus was an octave-band (2000-4000 Hz) of noise presented for four hours at 120 dB SPL.

### Cochlear Histological Analysis

Comprehensive histological analysis of this macaque cohort has been recently published (Mondul et al., 2026). We provide a brief overview here. Following completion of the current study, monkeys were euthanized and transcardially perfused. Brain and cochlear tissue was harvested for dissection and immunohistochemistry. Immunolabeling and confocal imaging of cochlear whole mounts was conducted to quantify IHC and OHC counts, IHC and OHC ribbon counts and sizes, and efferent terminal densities (Valero et al., 2017), and the methods and results described extensively in Mondul et al. (2026). Data from noise-exposed subjects were compared to unexposed subjects to assess anatomical integrity along the cochlear length.

### Psychometric and neurometric data analysis

Detection theoretic methods were used to analyze behavioral and neuronal responses, as well as simulated data from a model (Macmillan and Creelman, 2004; Swets, 1973; Tanner and Swets, 1954). Behavioral data were analyzed initially in terms of *d’* to facilitate comparison with O’Connor et al. (1999) and were subsequently analyzed in terms of probability correct to facilitate comparison with neuronal responses, detailed below.

Behavioral performance from each block of data was analyzed to calculate hit rate at each tone level (*H(level)*) and false alarm rate (*F*). Sensitivity was calculated as *d*^!^(*level*) = (*z*(*H*(*level*)) − *z*(*F*)), where *z* represents calculation of the z-score of the value, implemented in MATLAB via the function “norminv.” From *d’*, probability correct in a two-alternative forced-choice experiment was given by *pc(level)*, as *pc*(*level*) = *z*^−1^(*d*’(*level*)/2), where *z^-1^* represents the transformation from a standard normal variate to probability correct. Calculation of *pc,* rather than a more common *d’* measure, permitted comparison with our distribution free receiver operating curve (ROC) calculation of probability correct based on neuronal responses (Rocchi and Ramachandran, 2020, 2018). Psychometric and neurometric functions were fit with a modified Weibull cumulative distribution function (CDF) as others have done in both detection and discrimination tasks (Britten et al., 1996; Christison-Lagay and Cohen, 2014; O’Connor et al., 1999). The modified equation was: *pc*(*level*) = *c* − *d* ∗ *e*^−(*level*/*λ*)^***^κ^*** *for level* ≥ 0, where *c* represents saturation and *d* represents the range of the function, and *λ* and *κ* represent the threshold and slope parameters respectively. Often levels were presented that were below zero, and in these cases the Weibull fit was translated to higher levels before fitting, and translated back to the original levels after fitting. The threshold was calculated as that tone level at which *pc_fit_(level)*=0.76, and in the cases where *d’* was used, threshold criterion was 1.5 (after O’Connor et al., 1999).

Reaction times (RT) were calculated for all hit trials. They were calculated as follows: RT = Time of Lever Release – Time of Tone Onset. RTs as a function of tone level were fit with a line to provide RT slope and intercept, as described in previous studies from this laboratory (Dylla et al., 2013; Rocchi and Ramachandran, 2018).

The dynamic range (DR) of each psychometric function was calculated as the range of tone levels over which *pc_fit_(level)* spanned, from saturation minus 90% of that range, to saturation minus 10% of that range. This was done by using the *c* parameter as an estimate of saturated performance, and *d* as the total range, or amplitude of the psychometric function.

### Statistical analyses and curve-fitting

In all cases, curve fits were attained via non-linear least squares method implemented in MATLAB. Bayesian information criterion was calculated in MATLAB (2018a; Mathworks Inc.) using the non-linear model fit function, “fitnlm,” which returns multiple information criteria (including BIC), as well as goodness of fit measures R-squared and p-values. Threshold vs. duration trends were fit with an exponential function (Equation 1). All time constants (τ, the rate parameter), exponents (*m*, the rate parameter) and constants of proportionality (*I_k_* the range parameter) reported were taken from significant fits to the data (p < 0.05). The exponential function can be expressed as

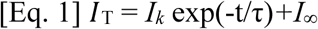

Statistical analysis of the effects of tone frequency, noise masker, and duration were conducted using linear mixed effects models (“fitlme”) in MATLAB 2018a. This allowed us to accommodate datasets with missing points, a common reason for avoiding repeated measures ANOVA in cases such as these (Krueger and Tian, 2004). The dependent variable in the models assessing the effects of tone frequency or noise masker was either τ or *I_k_*, the rate and range parameters, respectively. Background noise, tone frequency, and an interaction term between the two were entered as fixed effects into the model, while intercepts for individual macaques were entered as random effects. The effects of duration on threshold and DR were constructed by entering tone duration as a fixed effect, and the individual monkey as a random intercept term. Detection threshold or DR were entered as dependent variables. In all cases p-values were obtained by likelihood ratio testing of the model with the effect in question against the model without the effect in question. 2 sample Kolmogorov-Smirnov tests were used to assess differences between time constant distributions between brain structures, and between quiet and noise conditions (“kstest2” in MATLAB). Finally, correlations between noise-induced dynamic range increases and two measures of cochlear histopathology established in our previous study (Mondul et al. 2026); namely, relative inner hair cell synapse loss (normalized within subject) and inner hair cell ribbon synapse volume difference (median relative to a control group. Correlation analyses utilized the repeated-measures correlation (Bakdash and Marusich, 2017), which estimates the common within-animal association after removing between-animal differences, while accounting for the dependence between the ears. Similarly, the mixed effects model approach also allowed modeling between-animal differences with a random intercept term.

### Time window analysis

Neuronal responses were analyzed using different time windows, to facilitate comparison with the tone durations used in the behavioral experiments. The latency of the response to a 200 ms tone 5 dB above detection threshold was estimated using the point at which the cumulative spike function significantly deviated from a 250 ms baseline period (Rowland et al., 2007; Rowland and Stein, 2007). Starting at the latency, responses were then calculated using time windows of 100, 50, 25, 12.5, 6.5, and 3.25 ms. Using each of these time-windows, neurometric probability correct could be calculated at each tone level, using previously mentioned ROC analysis. Assessing how neurometric performance (e.g. threshold) changed as a function of time/duration provides a measure of temporal integration analogous to behavior when fit with a three-term exponential function (Eqn. 1).

## RESULTS

### Hierarchical differences in temporal integration emerge in masking noise

Tone evoked responses from the cochlear nucleus (CN) and inferior colliculus (IC) of macaques performing a tone detection task were analyzed using a time window analysis (see **Time window analysis** in the Methods section). These recordings are part of a previously published dataset (Mackey et al. 2023). From the tone evoked response (Figure 1A), the latency was calculated (Figure 1B), and neurometric accuracy (Probability Correct, see Methods) was then calculated via ROC analysis as a function of time elapsing after neuronal latency. This time after latency will be referred to as duration. Neurometric functions resembled psychometric functions: threshold and slope changed as a function of duration (Figure 1C).

**Figure 1.**
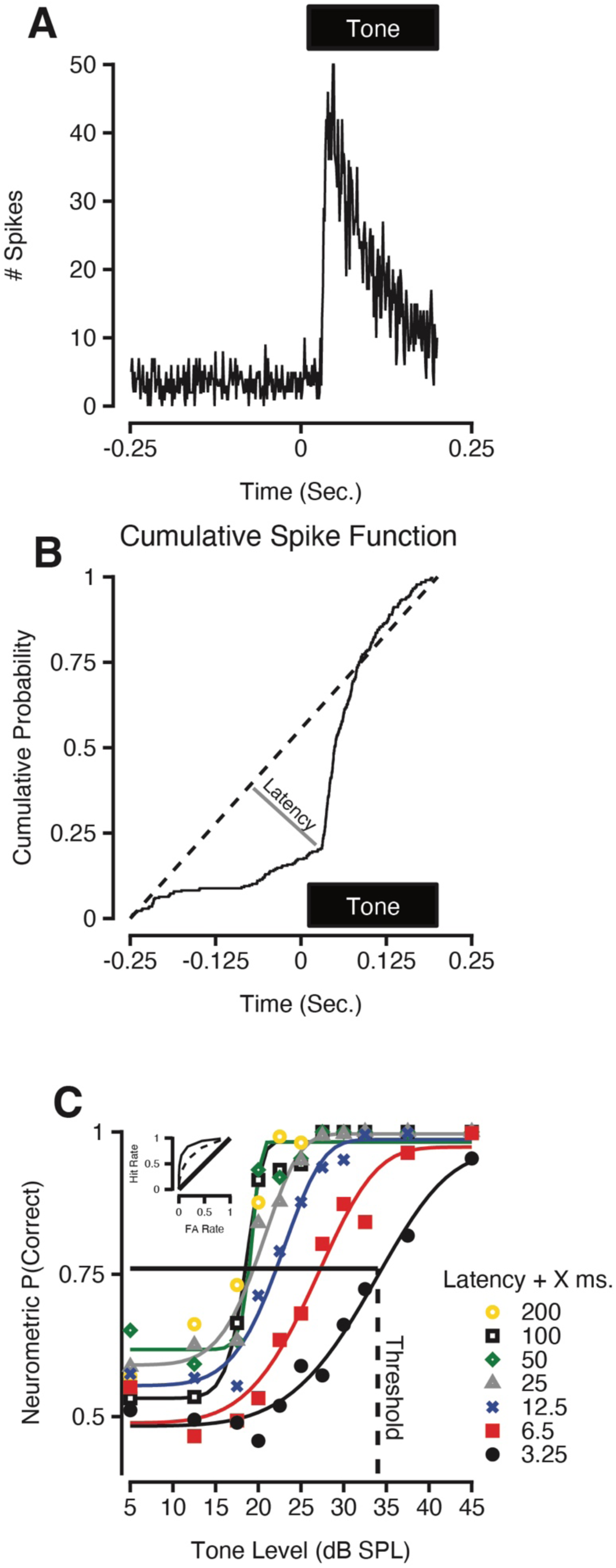
Time window analysis on single-unit responses. **A.** Example peristimulus time histogram from a neuron in the IC. **B.** Example cumulative spike function (CSF) based on the response in A. The CSF was used to estimate neuronal latency. **C.** Example neurometric functions calculated via ROC analysis (inset) at different durations, as indicated by the symbols in the legend. The solid line represents P(correct) = 0.76; threshold was the tone level at which the Weibull fit for PC as a function of level was 0.76.

Neurometric thresholds from each neuron were plotted as a function of duration and fit with an exponential function (Figure 2), to compare neurometric and psychometric indices of temporal integration such as time constants, which are inversely related to the rate of temporal integration. The effect of duration was consistent across the CN and IC, and the trend resembled behavioral trends in that threshold decreased exponentially as a function of duration. Figure 2A-B illustrates how thresholds were fit with exponential functions to extract the neurometric time constant, and representative neurons are shown. While the form of the exponential function (and thus the time constant) did not differ between quiet and noise conditions for the CN, an increase in the time constant by masking noise was regularly observed in the IC population (Figure 2C & D). Figure 2C-D shows median neurometric thresholds in quiet and in noise, in both brain regions, normalized to the lowest threshold (200 ms); i.e. each threshold by duration trend was normalized before taking the median across the neuronal population. . Hierarchical differences (between the IC and CN) in temporal integration were not apparent in quiet conditions, as can be seen in the green time constant distributions, plotted as cumulative probability distributions in Figure 2E and F. The CN and IC distributions were not significantly different in quiet, which was confirmed with a 2-sample Kolmogorov-Smirnov test (p = 0.29). However, in noise, a slower subpopulation emerged in the IC, which more closely approximated behavioral time constants, as can be seen in Figure 2F. This effect of brain region was validated with a two sample Kolmogorov-Smirnov test (*p* = 0.022), and the effect of noise in the IC was significant after correction for multiple comparisons (*p* = 0.017).

**Figure 2.**
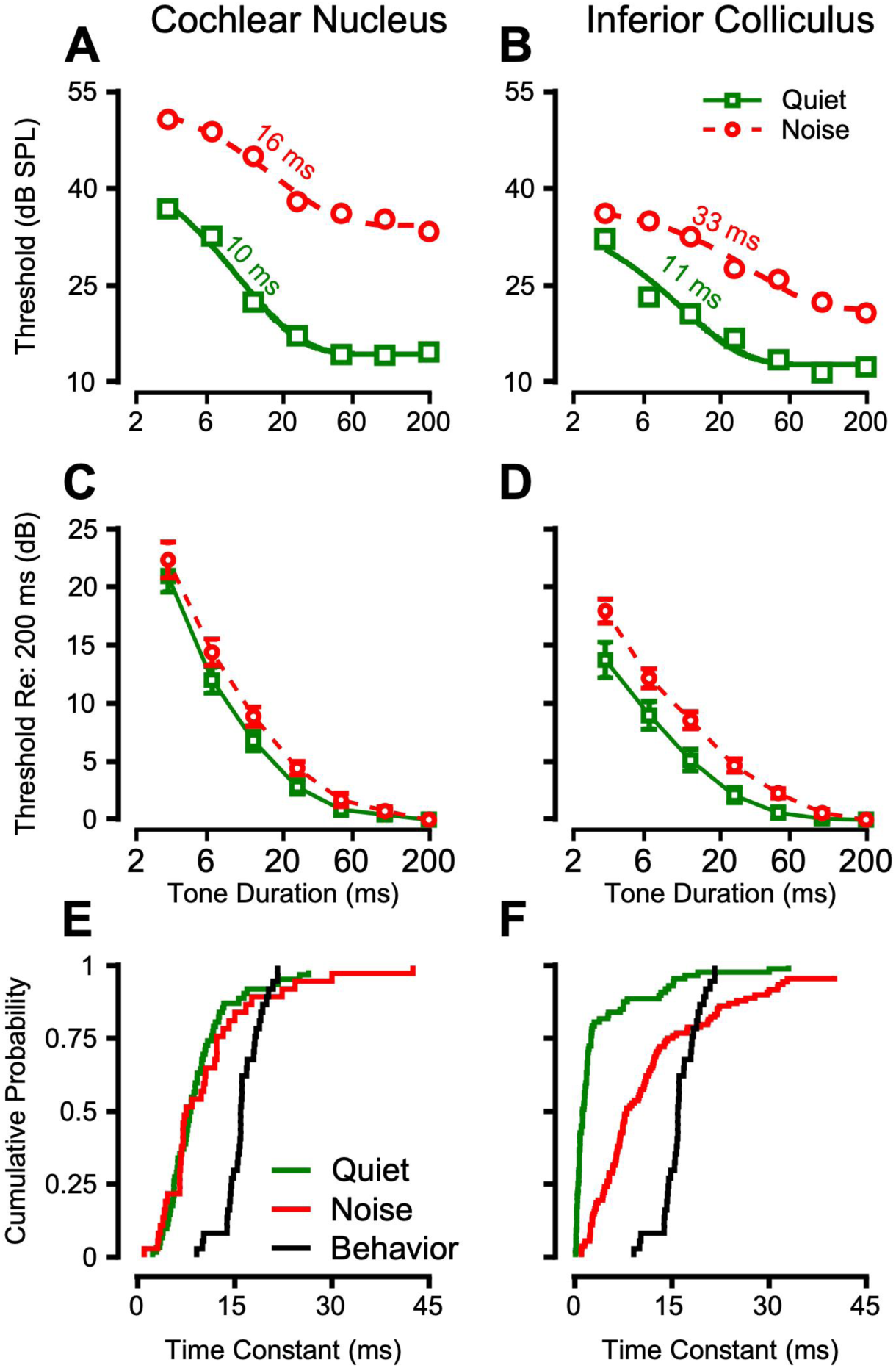
Neurometric measures of temporal integration reveal a hierarchical effect of noise. **A)** Example neurometric thresholds from the same neuron in quiet and in noise, fit with three term exponential functions. Temporal integration time constants are listed above each trace for the CN (**A**) and the IC (**B**). **C)** Median (+/- standard error) neurometric thresholds in the CN as a function of duration, in quiet (green squares) and in noise (red circles). **D)** Median neurometric thresholds in the IC. **E)** Time constants of all neurons in quiet (green) and noise (red) plotted cumulatively for the CN (**E**) and IC (**F**), along with the distribution of behavioral time constants (black). The effect of noise on time constants was significant in the IC as assessed by a 2-sample Kolmogorov-Smirnov test after Benjamini-Hochberg correction (*p* = 0.017). Behavioral time constants were previously published (Mackey et al. 2021).

### Effects of noise exposure on psychometric function dynamic range

We tested the hypothesis that psychophysical measures of temporal integration would be impaired by noise exposure that causes temporary threshold shifts (TTS) in male and female macaques. Detection of tones of varying duration was assessed in quiet both before and after exposure to a 120 dB SPL octave band noise (2-4 kHz) for four hours. Cochlear histological analysis indicates that this noise exposure causes synaptic hypertrophy throughout the cochlea, with little to no synapse loss (Mondul et al., 2026) (Figure 3B). Additionally, the TTS lasted less than two months in a large cohort of macaques (Figure 3A; Mondul et al., 2026). However, 4-7 months post exposure, psychometric function slopes to tones in quiet exhibited reliable changes at durations of 6-25 ms (Figure 4). Tone detection accuracy was quantified using psychometric dynamic range, as opposed to slope, to make clear whether the numerator or denominator of the slope was changing (DR; Figure 4). Perhaps somewhat counterintuitively, the changes to DR were concentrated at 1 kHz, distant from the 2-4 kHz band of the noise exposure.

**Figure 3.**
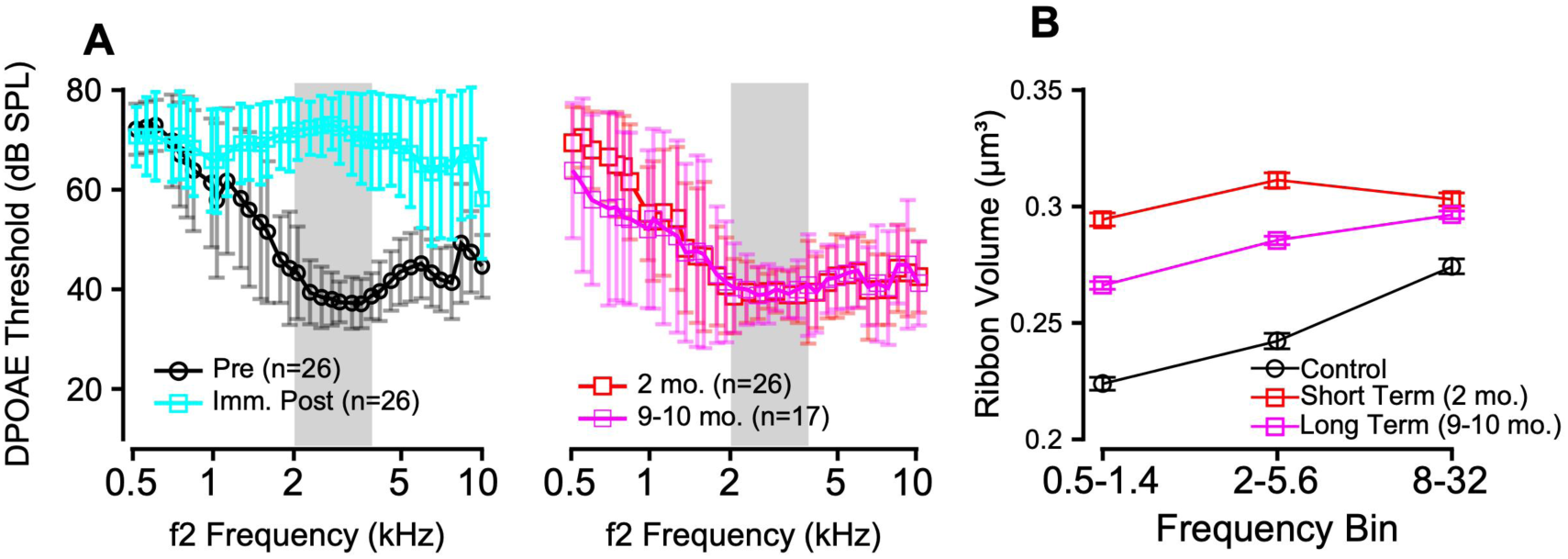
Noise-induced temporary threshold shift. **A**) Distortion product otoacoustic emissions (DPOAE) thresholds before (black), immediately post-, and at subsequent time-points following noise exposure confirmed the temporary depression of outer hair function. **B)** Inner hair cell ribbon volumes (Mean +/- SEM) in control and noise exposed NHPs.

**Figure 4.**
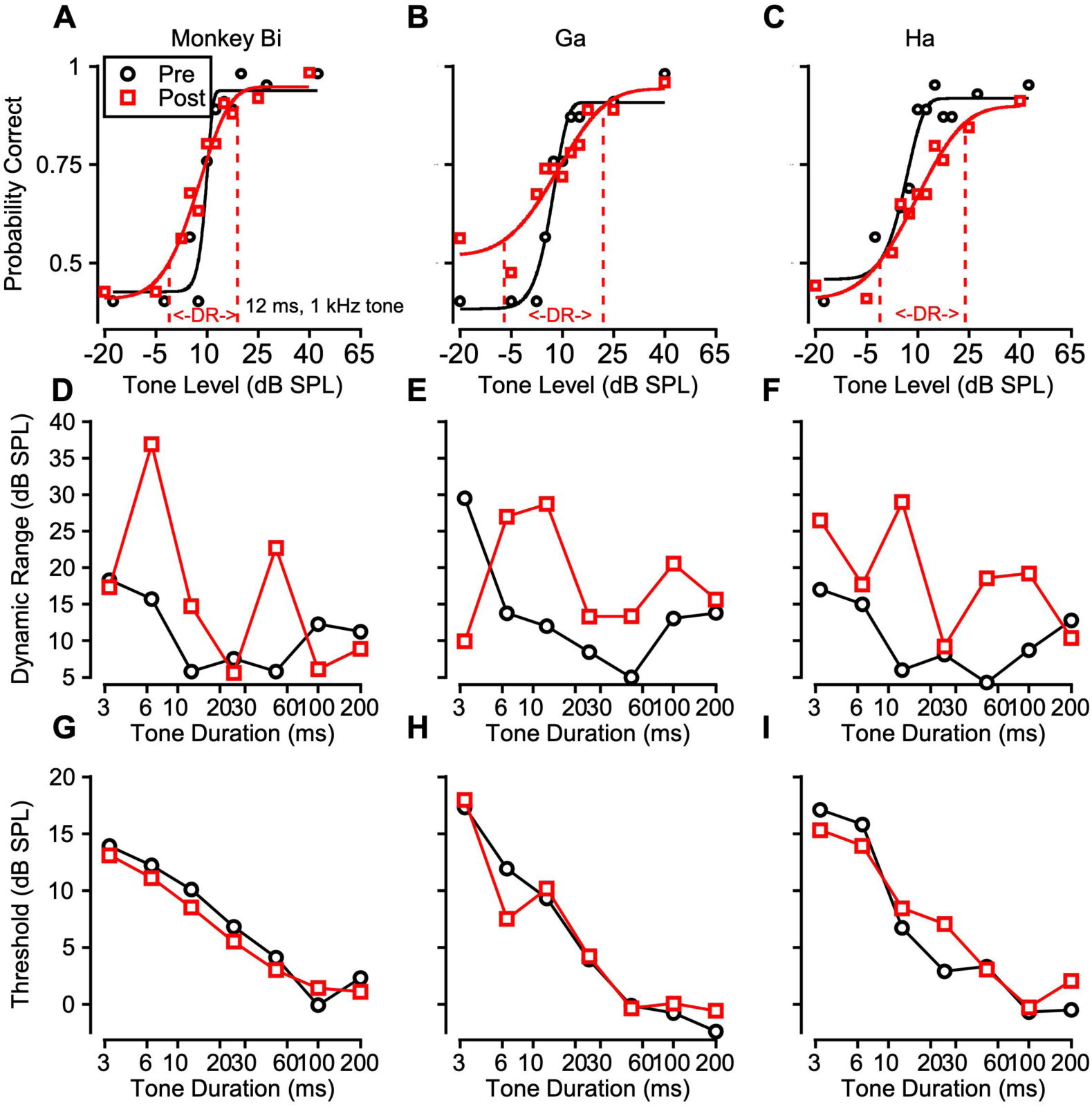
Change in dynamic range following noise exposure in a male macaque cohort. **A)** Psychometric functions from Monkeys Bi (**A**), Ga (**B**), and Ha (**C**) detecting 12 ms, 1 kHz tones in quiet, pre (black circles) and post (red squares) exposure. Dynamic range is indicated by the dashed red traces on each function. **D)** Dynamic range of 1 kHz psychometric functions as a function of tone duration pre and post exposure for each monkey (**D, E, F** for Monkeys Bi, Ga, and Ha, respectively). **G)** Detection threshold for 1 kHz tones in quiet, as a function of tone duration for monkeys Bi (**G**), Ga (**H**), and Ha (**I**).

In a mixed effects model incorporating both male and female data (Table 1), effects of noise exposure on psychometric function DR were significant, as well as interactions of noise exposure and sex, reflecting the fact that noise exposure impaired performance in both cohorts, but did so differently across frequency and duration. Detection thresholds (*PC =* 0.76; Figure 4) were unchanged after noise exposure in male macaques (Figure 4 G-I). Mixed effects modeling of the male data alone revealed a significant effect of noise exposure on DR, as well as interactions of exposure with tone frequency and duration. Female macaques showed similar effects, with a significant effect of noise exposure, as well as an interaction with tone duration (Figure 5; Table 1). The magnitude of the noise exposure effect was substantial, with the joint male/female model predicting an 8.1 dB increase in DR after noise exposure, which in many cases was a doubling in DR (Table 1; Figure 4; Figure 5 Top Row). The effect of sex was that DR was larger in the female cohort at baseline, indicated by a positive t-statistic, and a model coefficient estimate of 3 dB (Table 1). Another key difference in the male and female cohorts was that thresholds were elevated after noise exposure in female monkeys Pi and Lu by 8-15 dB at short durations (3-25 ms), which can be seen in Figure 5 (Bottom row). Surprisingly, this effect was not restricted to the tone frequency at which DR was elevated. Thus, thresholds at different frequencies are shown in Figure 5 (bottom row) for the female cohort, where occasional permanent threshold shifts, restricted to short durations, were observed.

**Figure 5.**
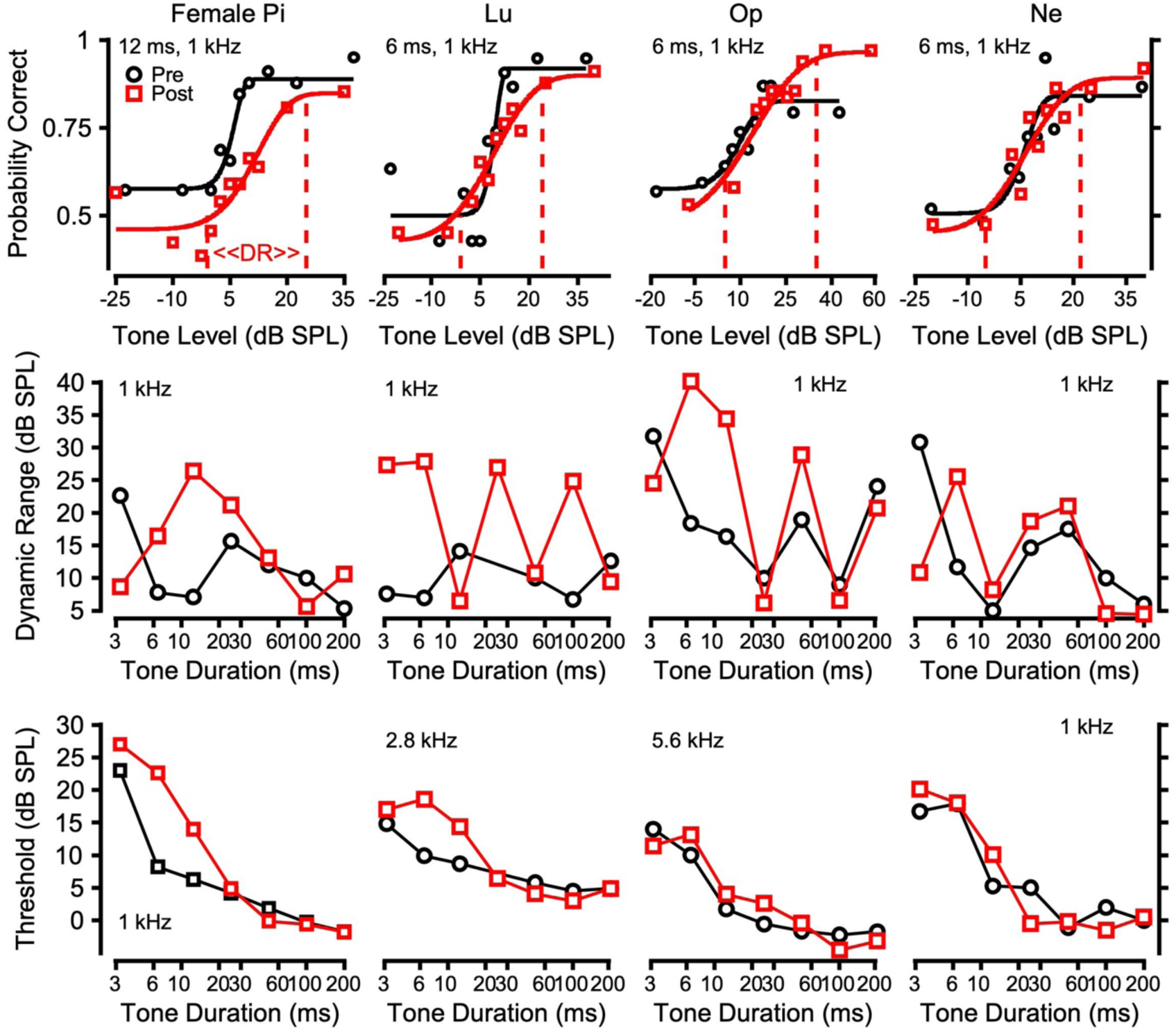
Change in dynamic range and threshold following noise exposure in a female cohort. **Top row,** examples of DR change following noise exposure. Psychometric functions from monkeys Pi, Lu, Op, and Ne are shown. Formatting follows conventions in the last figure. **Middle row,** DR as a function of duration at 1 kHz for each monkey. **Bottom row,** detection threshold as a function of tone duration for each monkey at select frequencies exhibiting permanent threshold shifts at short durations only.

**TABLE 1.** Mixed effects model analysis of the effects of noise exposure and sex on psychometric function dynamic range (DR).

| **<u>MODEL</u>** | **<u>Effect of noise exposure (t-stat, p-value)</u>** | **<u>Effect of sex (t-stat, p-value)</u>** |
| --- | --- | --- |
| **<u>Male cohort</u>**<br>Psychometric DR ~ Noise<br>Exposure*Tone Frequency + Noise<br>Exposure*Duration + Tone<br>Frequency + Duration + Noise<br>Exposure + (1\|Monkey) | Noise Exposure: $t = 1.23, p = 0.0019$<br>Noise Exp*Tone Freq: $t = -2.3, p = 0.019$<br>Noise Exp*Duration: $t = 2.26, p = 0.024$ | N/A |
| **<u>Female cohort</u>**<br>Psychometric DR ~ Duration* Tone<br>Frequency*Noise Exposure Status +<br>Duration + Tone Freq + Noise<br>Exposure Status + (1\|Monkey) | Noise exposure: $t = 2.4, p = 0.01$<br>Noise Exp*Duration: $t = -2.1, p = 0.039$ | N/A |
| **<u>Male & Female</u>**<br>Psychometric DR ~ Noise<br>Exposure*Sex*Tone Frequency +<br>Noise Exposure + Sex + Tone Freq +<br>Duration + (1\|Monkey) | Noise exposure: $t = 3.28, p = 0.001$ | Sex: $t = 2.8, p = 0.005$<br>Noise Exposure*Sex = $t = -2.12, p = 0.034$ |

### Effects of noise exposure on reaction time

The effects of noise exposure on reaction times were also assessed. Previous work has characterized how RTs increase with sound level, and signal-to-noise ratio (Dylla et al., 2013; Kemp, 1984; Kemp and Irwin, 1979). These effects have been quantified using a linear fit to RT as a function of tone level (Bohlen et al., 2014; Dylla et al., 2013; Rocchi and Ramachandran, 2018). The two parameters of the RT vs. level function (slope and intercept) were used in the present analysis to assess two potential effects of noise exposure on RTs: a change in the growth of loudness (slope change), or an overall change in the speed of auditory processing (intercept change). As can be seen in Figure 6A, RT slope and intercept changed jointly following noise exposure, with slope being reduced, and intercept decreasing. The intercepts and slopes of the reaction time vs. tone level linear regressions were entered into a mixed effects model as the response variable, with tone frequency, sex, duration, and pre vs. post noise exposure status as fixed effects. Individual monkeys were entered into the model as random effects. Reaction time intercept was lower after noise exposure, indicating faster RTs, possibly a result of training (Coefficient estimate: -29.97 ms, t = -4.17, p = 4*10^-5^). Reaction time slope was reduced at short tone durations following noise exposure, indicated by a significant positive interaction between noise exposure status and tone duration (t = 2.0, p = 0.043). At 3.25 and 6.5 ms, intercepts were decreased and slopes were reduced (Figure 6B, Figure 6C) at the group level, likely driving the changes in the mixed effects model analysis. However, as can be seen in Figure 6B and Figure 6C, trends were variable across duration and should be interpreted conservatively.

**Figure 6.**
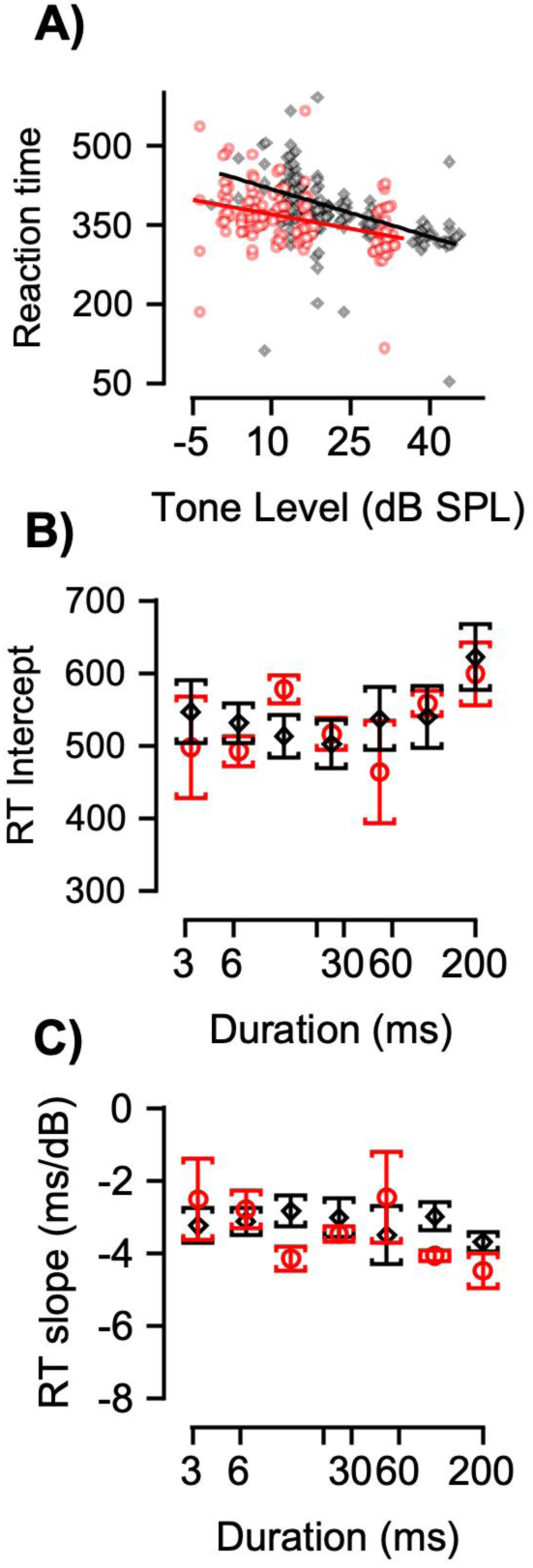
Change in reaction-time following noise exposure. **A)** Example reaction times before (black) and after (red) noise exposure from Monkey Bi, detecting 6.5 ms, 1 kHz tones in quiet. Lines are linear regression. **B)** Mean intercept from RT vs. tone level regressions across all macaques RTs to 1 kHz tones. **C)** Mean slopes from RT vs. tone level regressions across all macaques RTs to 1 kHz tones. Error bars are standard error of the mean in all cases.

We also conducted all of the behavioral experiments described here in the presence of continuous, 76 dB SPL background noise, but found no deficits due to noise exposure. Thresholds (p = 0.10) and reaction time measures (p > 0.05) from the masked detection experiments did not significantly differ before and after noise exposure as assessed by mixed effects model analysis. The dynamic range of the psychometric function slightly *decreased* following noise exposure, possibly another beneficial effect of training (t = -0.23, p = 0.004).

We hypothesized that variability in noise-induced perceptual deficits would correlate with variability in cochlear histopathology. Two primary metrics emerged from the previous analysis of cochlear histopathology: inner hair cell (IHC) ribbon synapse counts and IHC presynaptic ribbon volumes (Mondul et al. 2026). To assess the correlations, we utilized repeated measures correlation (Bakdash and Marusich, 2017) and a linear mixed effects model which allowed us to account for the dependence between the ears, and the repeated measures within NHP. Correlations were performed with normalized synapse counts or median ribbon volume differences against within-subject psychometric function dynamic range at corresponding frequencies and varying durations. Neither approach suggested correlations between cochlear histopathology and changes in psychometric function dynamic range, (Table 2). The trend towards significance for IHC synapse volume difference suggested an inverse correlation (i.e. greater hypertrophy correlated with better perception following noise exposure), which is inconsistent with the main finding (noise exposure disrupts perception) and thus merits further consideration in future studies before it can be meaningfully interpreted.

**Table 2.** No correlation between noise-induced dynamic range increase and cochlear histopathology.

| **<u>Predictor</u>** | **<u>RM corr R-value</u>** | **<u>RM corr R-value CI</u>** | **<u>RM corr P-value</u>** | **<u>LME beta</u>** | **<u>LME P-value</u>** |
| --- | --- | --- | --- | --- | --- |
| IHC Rel Syn Count | 0.084 | [0.02, 0.16] | 0.533 | 6.4 | 0.335 |
| IHC Syn Vol Diff | -0.215 | [-0.35, 0.02] | 0.077 | -20.13 | 0.061 |

## Discussion

These results contribute in many ways to the understanding of how acoustic evidence is accumulated over time. Analysis at the behavioral, neurophysiological, and computational levels indicate that 1) macaques are well-positioned to model human temporal integration (Mackey et al., 2021, 2024), 2) there is a hierarchical effect of noise on temporal integration in the subcortical auditory system, and 3) hidden hearing loss may be in part due to noise-induced synaptic dysfunction, and subsequent reduction in the rate of temporal integration.

### A hierarchical effect of noise on subcortical auditory temporal integration

Conclusions about the contribution of the CN and IC to temporal integration based on previous studies were complicated by the use of anesthetized animal preparations, lack of studies in nonhuman primates, and most studies measuring responses to stimuli presented in unnaturally quiet environments. Anesthetized/passive-listening preparations are limited because task engagement affects neuronal encoding along the auditory pathway (Downer et al., 2015; Niwa et al., 2012; Rocchi and Ramachandran, 2020; Ryan and Miller, 1977; Slee and David, 2015). The exclusive use of rodents and cats in previous studies begs the question of how these results translate to humans, because primate auditory systems differ from rodents and cats in anatomical organization (Hackett, 2011; Hackett et al., 2014; Kavanagh Moore, 1980), neurophysiological measures of temporal coding (Hoglen et al., 2018), auditory nerve frequency tuning (Joris et al., 2011; Verschooten et al., 2018), and perceptual measures of temporal processing (Kelly et al., 2006; Mackey et al., 2021, 2024; Moody, 1994). Finally, only presenting stimuli in quiet begs the question of how these results generalize to natural environments. Regarding the relevant brain regions in these studies, previous studies have posited that performance at short durations requires integration of populations of ANF inputs (Lopez-Poveda and Barrios, 2013; Mackey et al., 2021, 2024; Stevens and Wickesberg, 1999), a requirement that, at the anatomical level, is met by CN and IC. Building on these studies, and the Poisson process model in our previous study (Mackey et al., 2021), the present results investigated temporal integration in the CN and IC. In contrast to a previous study of anesthetized chinchilla CN (Clock et al., 1993), the CN time constants presented here did not resemble behavior in quiet or in noise. It was hypothesized that temporal integration in the IC would resemble behavioral temporal integration, as IC responses provide reliable estimates of psychometric threshold and slope, and exhibit significant choice-probabilities (Mackey et al., 2023). These neurometric-psychometric correlations may be due to the high degree of convergence in the midbrain, and consequent reduced variability of neuronal responses we previously reported (Mackey et al., 2023). However, IC temporal integration time constants only resembled behavioral time constants in the presence of noise. This wider range of timescales at which neurons encode sound may be of utility in noisy, suprathreshold environments, and/or for the encoding of complex sounds that exhibit fluctuation on multiple timescales (e.g. species-specific communication sounds). The hierarchical differences between CN and IC are consistent with a study of IC, thalamus, and primary auditory cortex (Asokan et al., 2021), which characterized how progressively longer timescales are encoded in these regions. Asokan et al. did not explicitly relate neural measures to behavioral measures, but the CN and IC results presented here may suggest that auditory cortex time constants would more closely resemble behavioral measures of temporal integration. Further downstream, parietal cortex integrates acoustic evidence (Yao et al., 2023, 2020), and sensory evidence more generally (Cohen et al., 2004; Huk and Shadlen, 2005). Alternatively, the superior colliculus can integrate acoustic evidence, as multiple studies indicate (Jay and Sparks, 1987; Rajala et al., 2017; Wallace et al., 1996), and there is causal evidence for its role in evidence accumulation in the visual domain (Jun et al., 2021). Future studies can empirically address how neural measures of temporal integration in these regions relate to perceptual measures.

### Effects of noise exposure and cochlear pathology

Our NHP model documents long-lasting perceptual and neurophysiological changes after a noise exposure that induced TTS and persistent synaptic hypertrophy without significant synapse loss (Figure 3; Mackey et al., 2026; Mondul et al., 2026). The present results pose foundational questions about the relationship between hair cell synaptic function and temporal integration. How can synaptic hypertrophy, in the absence of overt synaptopathy, alter temporal integration? Synaptic hypertrophy has been linked to glutamatergic excitotoxicity and altered release dynamics; specifically, studies have noted ultrastructural evidence of reduced mitochondrial content, swollen endoplasmic reticulum, and longer response latency preceding overt synaptopathy (Pal et al., 2025; Sheets et al., 2017; Robertson, 1983). There are heterogeneous effects across auditory nerve fiber populations, which are differentially susceptible to excitotoxicity, and altered glutamatergic signaling (Moverman et al., 2023; Reijntjes et al., 2026; Suthakar and Liberman, 2022; Vincent et al., 2024). Subsequent to these changes in glutamatergic signaling, auditory nerve fiber demyelination likely contributes to post-noise perceptual and neurophysiological deficits by degrading conduction velocity and temporal precision (Kohrman et al., 2020; Wan and Corfas, 2017; Zhang et al., 2025). Across successive stages in the auditory pathway, reduced precision of signal conduction, and thus, postsynaptic temporal integration, could be magnified. This conclusion is consistent with the Poisson model introduced in our previous study, which suggested reduced auditory nerve fiber recruitment can result in increased psychometric DR (Mackey et al., 2021). Moreover, this is corroborated by modeling studies predicting that deafferentation should alter the perception of short duration stimuli (Lopez-Poveda, 2014; Lopez-Poveda and Barrios, 2013; Mackey et al., 2021; Stevens and Wickesberg, 1999).

Previous studies of have investigated temporal integration as a candidate behavioral assay of cochlear synaptopathy (Marmel et al., 2020; Wong et al., 2019) and reported null results. Here it was shown that temporal integration was impaired by noise exposure that causes IHC synapse hypertrophy. The discrepancy between these studies may be due to differences in psychometric methodologies or synaptopathic pathology. Marmel et al. (2020) and Wong et al. (2019) measured threshold using an adaptive procedure, and psychometric function slope/DR are not routinely calculated in such studies. It may be that subjects in these studies had perceptual deficits not evident in detection threshold, as was the case for many of the monkeys in this study. Another difference is that Wong et al. used a glutamate analog, kainic acid, to mimic the effects of synaptopathic noise exposure. Kainic acid causes substantial synaptic loss, whereas our noise exposure did not. These different manifestations of synaptic pathology may explain the discrepant results. Finally, Marmel et al. measured temporal integration in humans with *putative* synaptopathy. The lack of histological verification makes it unclear whether their null result was due to lack of synaptic pathology, or that synaptopathy doesn’t impair detection of short tones.

Interestingly, it was shown here that noise exposure’s effect on psychometric DR was not largest for the detection of 3.25 ms tones, but rather for 6-25 ms (see Results). While counterintuitive, this may be explained by a companion study from our lab, and from a study of mice auditory nerve responses. Specifically, (Suthakar and Liberman, 2022, 2021) found larger ANF onset responses and masked ABR Wave-I amplitudes in mice following synaptopathic noise exposure, and our cohort of noise-exposed NHPs displayed a similar increase in ABR Wave-I amplitudes to clicks in quiet following noise exposure (Mondul et al. 2026). In the present study, 3.25 ms tones were presented at relatively high sound levels to elicit detection behavior. Combined with their short duration (and consequentially broader spectra), these stimuli may serve to broadly recruit cells across the tonotopic map, with a high-frequency bias, in a similar fashion to clicks. It may be that 6-25 ms tones are in an ideal range to avoid enhanced click-like responses associated with shorter durations, while simultaneously lacking the robust responses elicited by longer (> 25 ms) duration tones. Future implementations of temporal integration as a biomarker of cochlear synaptopathy should carefully titrate stimulus parameters to optimize assay sensitivity.

A recent study found that kainic acid induced synaptopathy in budgerigars impaired detection in an “overshoot” paradigm, where tone and noise onset synchrony are manipulated to probe the perception of acoustic onsets (Loftus et al., 2026). Interestingly, normal hearing budgerigars don’t exhibit the overshoot effect commonly observed in humans and nonhuman primates (Rocchi et al., 2017), but Loftus et al. found that the perception of short tones at the onset of a noise masker was impaired following kainic acid administration. Loftus et al. speculated that this could lend insight into how synaptopathy impairs speech perception in humans, i.e. by altering the perception of brief transients that are critical for parsing phonemes. These findings, coupled with the present data indicate that investigation of acoustic onset perception is also likely to yield promising results in future studies of cochlear synaptopathy and hidden hearing loss.

## Acknowledgements

The authors would like to acknowledge Bruce and Roger Williams for fabrication of hardware, and Mary Feurtado for assistance with surgical procedures. The authors thank Dr. Amy Stahl, Dr. Jessica Feller, Namrata Temghare, Alejandro Tarabillo, and Jackson Mayfield for assistance collecting behavioral data. The authors thank Drs. M. Charles Liberman, Leslie Liberman, and Troy A. Hackett for conceptual input and assistance collecting histological data. The study was supported by research grant NIH R01 DC 015988 (MPIs R. Ramachandran and B. Shinn-Cunningham). CM was supported by The Department of Hearing and Speech Sciences, and the Ruth Kirchstein pre-doctoral fellowship from NIDCD (F31 DC 019823). JAM was supported by the Ruth Kirchstein fellowship for audiologists, from NIDCD (F32 DC 019817).

## Author Contributions

RR and CAM designed experiments. CAM, JAM, and RR conceptualized analysis. CAM and JAM collected data. CAM and JAM analyzed data. CAM wrote the paper. All authors revised and approved the final version of the paper.

